# Instant polarized light microscopy pi (IPOLπ) for quantitative imaging of collagen architecture and dynamics in ocular tissues

**DOI:** 10.1101/2023.01.29.526111

**Authors:** Po-Yi Lee, Hannah Schilpp, Nathan Naylor, Simon C. Watkins, Bin Yang, Ian A Sigal

## Abstract

Collagen architecture determines the biomechanical environment in the eye, and thus characterizing collagen fiber organization and biomechanics is essential to fully understand eye physiology and pathology. We recently introduced instant polarized light microscopy (IPOL) that encodes optically information about fiber orientation and retardance through a color snapshot. Although IPOL allows imaging collagen at the full acquisition speed of the camera, with excellent spatial and angular resolutions, a limitation is that the orientation-encoding color is cyclic every 90 degrees (π/2 radians). In consequence, two orthogonal fibers have the same color and therefore the same orientation when quantified by color-angle mapping. In this study, we demonstrate IPOLπ, a new variation of IPOL, in which the orientation-encoding color is cyclic every 180 degrees (π radians). Herein we present the fundamentals of IPOLπ, including a framework based on a Mueller-matrix formalism to characterize how fiber orientation and retardance determine the color. The improved quantitative capability of IPOLπ enables further study of essential biomechanical properties of collagen in ocular tissues, such as fiber anisotropy and crimp. We present a series of experimental calibrations and quantitative procedures to visualize and quantify ocular collagen orientation and microstructure in the optic nerve head, a region in the back of the eye. There are four important strengths of IPOLπ compared to IPOL. First, IPOLπ can distinguish the orientations of orthogonal collagen fibers via colors, whereas IPOL cannot. Second, IPOLπ requires a lower exposure time than IPOL, thus allowing faster imaging speed. Third, IPOLπ allows visualizing non-birefringent tissues and backgrounds from tissue absorption, whereas both appear dark in IPOL images. Fourth, IPOLπ is cheaper and less sensitive to imperfectly collimated light than IPOL. Altogether, the high spatial, angular, and temporal resolutions of IPOLπ enable a deeper insight into ocular biomechanics and eye physiology and pathology.

**Highlights:**

- We introduce IPOLπ, addressing IPOL limitations for characterizing eye collagen.
- IPOLπ orientation-encoded color cycle is 180° (π radians) instead of 90° in IPOL.
- IPOLπ requires a lower exposure time than IPOL, allowing faster imaging speed.
- IPOLπ visualizes non-birefringent tissues and backgrounds from brightness.
- IPOLπ is cheaper and less sensitive to imperfectly collimated light than IPOL.

## 1. Introduction

Collagen is a primary load-bearing component in ocular tissues. Its architecture determines the ocular biomechanical environment and susceptibility to several vision threatening conditions. (Coudrillier et al., 2012; Ethier et al., 2004) Therefore, characterizing collagen fiber organization and biomechanics is important to fully understand eye physiology and preventing vision loss.

Polarized light microscopy (PLM) has been used to study collagen architecture in soft tissues for several decades. (Canham et al., 1991; Diamant et al., 1972; Keefe et al., 1997; Tower et al., 2002) A major advantage of PLM over other more conventional histology is that it does not require labels or stains, avoiding potential artifacts and simplifying preparation. (Koike-Tani et al., 2015) Over the past few years widefield PLM has continued to be refined and recently demonstrated robust for characterizing collagen architecture of posterior pole ocular tissues. (Brazile et al., 2018; Filas et al., 2014; Gogola et al., 2018a; Gogola et al., 2018b; Jan et al., 2018; Jan et al., 2017a; Jan and Sigal, 2018; Jan et al., 2015b; Jan et al., 2017c; Yang et al., 2018a; Yang et al., 2018b) Other imaging techniques leveraging polarized light have also been used extensively to study ocular collagen, including polarization sensitive second harmonic generation (PS-SHG) (Cisek et al., 2021; Gusachenko et al., 2012; Mansfield et al., 2008) and polarization sensitive optic coherent tomography (PS-OCT) (Baumann et al., 2014; Willemse et al., 2020; Yamanari et al., 2014). Nevertheless, widefield PLM has several strengths over raster-scanning techniques that continue to make it highly useful for soft tissues. PLM produces images with high, micrometer-scale, resolution over wide regions at high imaging speeds. PLM is substantially simpler and therefore cheaper. (Higgins, 2010; Whittaker and Przyklenk, 2009) The high angular resolution and sub-pixel information of PLM allows accurate measurement of small fiber undulations, or crimp. (Jan et al., 2015a; Kalwani et al., 2013) This is of particular interest for the study of the relationship between tissue architecture and biomechanics as it allows quantifying fiber crimp without the need to discern or trace individual fibers. Fiber tracing is very demanding on image resolution and analysis time, often leading to a reduced number of measurements. The above have made PLM a preferred technique for measuring collagen crimp.

Conventional quantitative PLM, however, requires multiple images at different polarization states to compute the collagen structure and orientation. (Shribak and Oldenbourg, 2003) Multiimage acquisition not only limits the imaging speed to evaluate the static architecture or quasistatic behavior in ocular tissues but may also introduce quantitative errors from post-processing among images, such as image registration.

Instant polarized light microscopy (IPOL) is a recently introduced technique that optically encodes information about fiber orientation and retardance through a color snapshot. (Lee et al., 2022; Yang et al., 2021) IPOL allows quantitative imaging of collagen at the full acquisition speed of the camera, with excellent spatial and angular resolutions. IPOL, however, has the limitation that the orientation-encoded colors are cyclic every 90 degrees (π/2 radians). In consequence, two orthogonal fibers have the same color and therefore the same orientation when quantified by color-angle mapping. Techniques to distinguish the orientations of orthogonal fibers have been suggested, (Keikhosravi et al., 2021) but the detailed methodology to quantify orientation and retardance have not been described.

Our goal in this work was to demonstrate IPOLπ, a new variation of IPOL, in which the orientation-encoding color is cyclic every 180 degrees (π radians). We describe how to use IPOLπ to conduct quantitative analysis on static and dynamics of ocular tissues, with high spatial and temporal resolutions. We present the fundamentals of IPOLπ, including a framework based on a Mueller-matrix formalism to characterize how fiber orientation and retardance determine the color. The improved quantitative capability of IPOLπ enables further study of essential biomechanical properties of collagen in ocular tissues, such as fiber anisotropy and crimp. We present a series of experimental calibrations and quantitative procedures to visualize and quantify ocular collagen orientation and microstructure in the optic nerve head, a region in the back of the eye.

## 2. Methods

This section is organized into five parts. In Section 2.1, we describe the configuration of IPOLπ imaging system. In Section 2.2, we introduce a framework based on a Mueller-matrix formalism to characterize how the color changes in fiber orientation and retardance. This simulation helps builds a foundation of quantitative analysis from the proposed imaging. In Section 2.3, we demonstrate how to experimentally calibrate the relationship between color and fiber orientation and the relationship between color and retardance. In Section 2.4, we introduce how to use the interpolating functions obtained from the experimental calibration in Section 2.3 to quantify images acquired by IPOLπ and then to visualize the quantitative results. Finally, in Section 2.5, we demonstrate the applications of IPOLπ for visualizing and quantifying static collagen architecture and dynamic collagen deformation under uniaxial stretch testing of optic nerve head tissues.

### 2.1 IPOLπ imaging system

The optic design of IPOLπ was to retrofit a commercial inverted microscope (Olympus IX83, Olympus, Tokyo, Japan) with a white light source, circular polarizer, polarization decoder, and color camera (DP74, Olympus, Tokyo, Japan) (**Figure 1**). Polarization decoder consists of a z-cut quartz and a linear polarizer. In the absence of a birefringent sample, the spectrum of the white light was unchanged and the image background appeared grey. With a birefringent sample, such as collagen, the spectrum of the white light was changed and thus the light appeared colorful. The frame rate of IPOLπ was limited only by that of the color camera, which in our setup was 60 frames per second.

**Figure 1.**
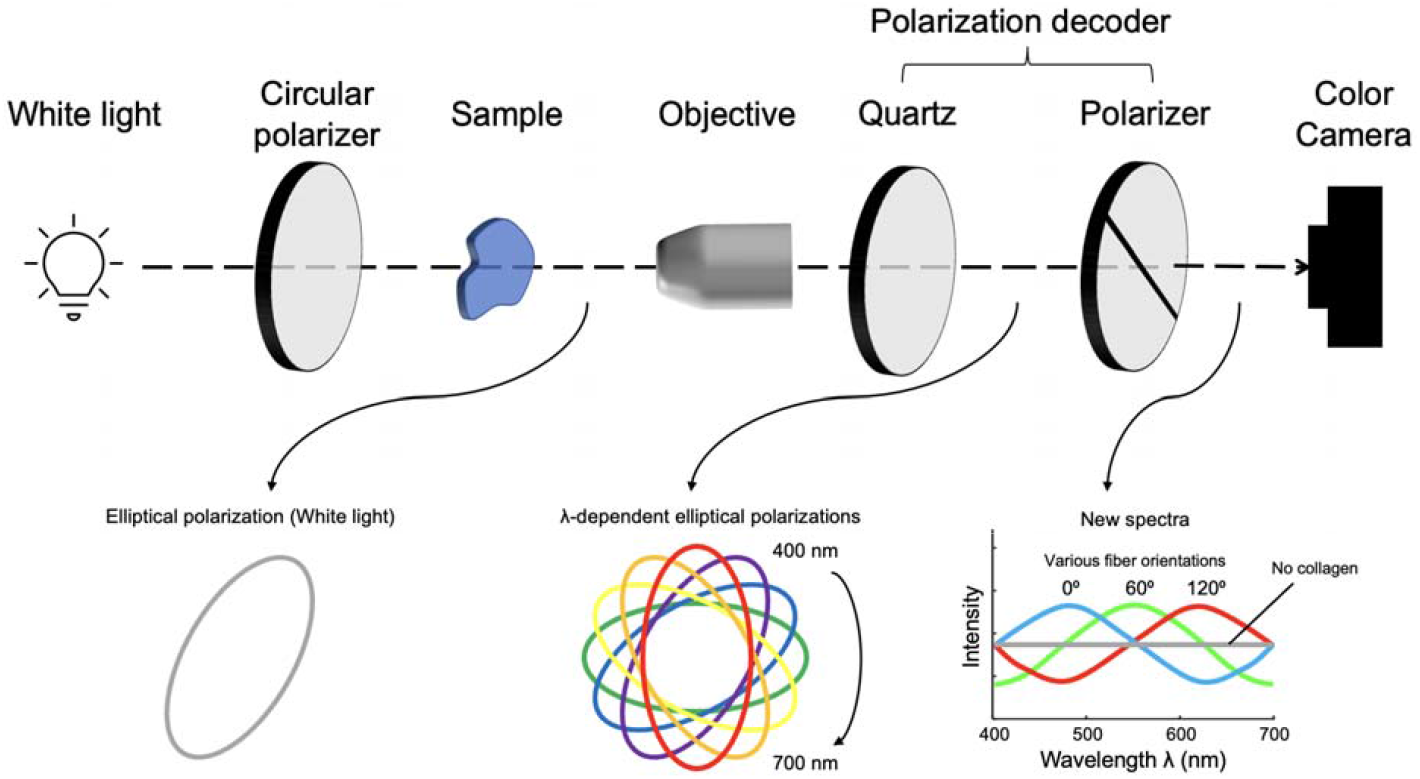
IPOLπ imaging configuration. A circular polarizer and a polarization decoder are retrofitted into the illumination and imaging paths, respectively. An alternative to the circular polarizer is to use a linear polarizer followed by a quarter-wave plate whose slow and fast axes are at 45°. The locations of the circular polarizer and the polarization decoder are swappable. The polarization of the white light is converted into an elliptically polarized light by passing it through the circular polarizer and the sample, where the orientation of the elliptical polarization depends on local fiber orientation. A new spectrum is then generated after the elliptically polarized light passed through the polarization decoder, and thus the output light is colorful. A color camera acquires the colorful light to produce true-color images indicating collagen fiber microstructure.

### 2.2 IPOLπ simulation

The polarization simulation was used to characterize the relationship between the output RGB color from IPOLπ and the material properties of collagen. We applied a Mueller-matrix formalism to simulate how the polarization states of the broadband white-light spectrum were altered through optical elements and a sample. In the Mueller-matrix formalism, briefly, a series of 4×4 transfer matrices, i.e., Mueller matrices, was introduced to operate on an incident 1×4 Stokes vector to obtain the corresponding transmitted Stokes vector. (Collett, 2005) The Stokes vector *S*_out_ of the output light from IPOLπ can be described as

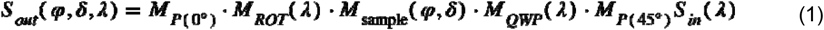

where M were the Mueller matrices and S were the Stokes vectors. The spectrum of the incident light *S*_in_ was referred to our LED white light source (IX3-LJLEDC, Olympus, Tokyo, Japan). A linear polarizer M_P_ orientated at 45 degrees and a quarter waveplate was equivalent to a circular polarizer. We simulated and compared the effects on the output light of four quarter waveplates: ideal (i.e., wavelength independent), achromatic (AQWP05M-580, Thorlabs, NJ, USA), zero-order (WPQ05ME-546, Thorlabs, NJ, USA), and multi-order (WPMQ05M-532, Thorlabs, NJ, USA). A phase retarder, M_sample_, was used to represent the birefringent sample, such as collagen fibers, where the slow axis orientation □ varying from 0 to 180 degrees, and the retardance δ varying from 0 to π radians. A z-cut quartz was model as a polarization rotator, M_ROT_, where the rotated angles were wavelength dependent. The polarization rotator allowed diverged the polarization directions of the visible spectrum (400-700 nm) within 180 degrees. The last one was a linear polarizer, M_P_, orientated at 0 degrees, also called an analyzer. For a given fiber orientation and retardance, S_out_ was calculated for the wavelengths between 400nm and 700 nm, where the first parameter of the 1×4 Stroke vector represented intensity. Each wavelength had a corresponding RGB (Red, Green, Blue) value, an additive color model. The RGB color was subsequently obtained by spectrally mixing the wavelength-dependent intensity of the output. The simulated color was converted from the RGB color space to the HSV (Hue, Saturation, Value) color space (Schanda, 2007). Note that the meanings of the terms “value” and “brightness” were the same in this study.

### 2.3 RGB color calibration

With the understanding of the relationship between color and material properties of collagen from the Section 2.2, we performed an experimental calibration and developed an algorithm to build system-specific color-angle and color-energy conversion maps. A chicken Achilles tendon was dissected and fixed with 10% formalin for 24 hours while under load. (Yang et al., 2018b; Yang et al., 2021) Following fixation, the tendon was cryo-sectioned longitudinally into 20-μm-thick sections. The tendon section fixed while under load was considered a uniform collagen organization with consistent fiber orientation. IPOLπ images were acquired with the chicken tendon section at several controlled angles relative to the longitudinal fiber direction, from 0 to 180 degrees, every 2 degrees.

Post-processing was necessary to extract the averaged color from the section and actual rotation angles due to regional variations in fiber content and orientation and the limited rotational accuracy. The individual images were registered and the rotation angles were obtained from the transformation matrices using Fiji software. (Schindelin et al., 2012) Then, with a region of interest manually placed on the color-uniform area in the stack, the RGB colors were extracted and averaged from the region of interest from the corresponding frames. The calibrated background was the mean of the colors obtained from all frames. The distance between color on experimental calibration and calibrated background as a unit of “energy”, which was related to retardance. We used the experimental calibration to generate pseudo calibrations with corresponding orientations and energies (**Figure 2**). The energy in low-brightness pseudo calibrations was weighted by the brightness to avoid image noise. We interpolated a combination of pseudo and experimental calibrations to build interpolating functions for orientation and energy inquiry. (Amidror, 2002)

**Figure 2.**
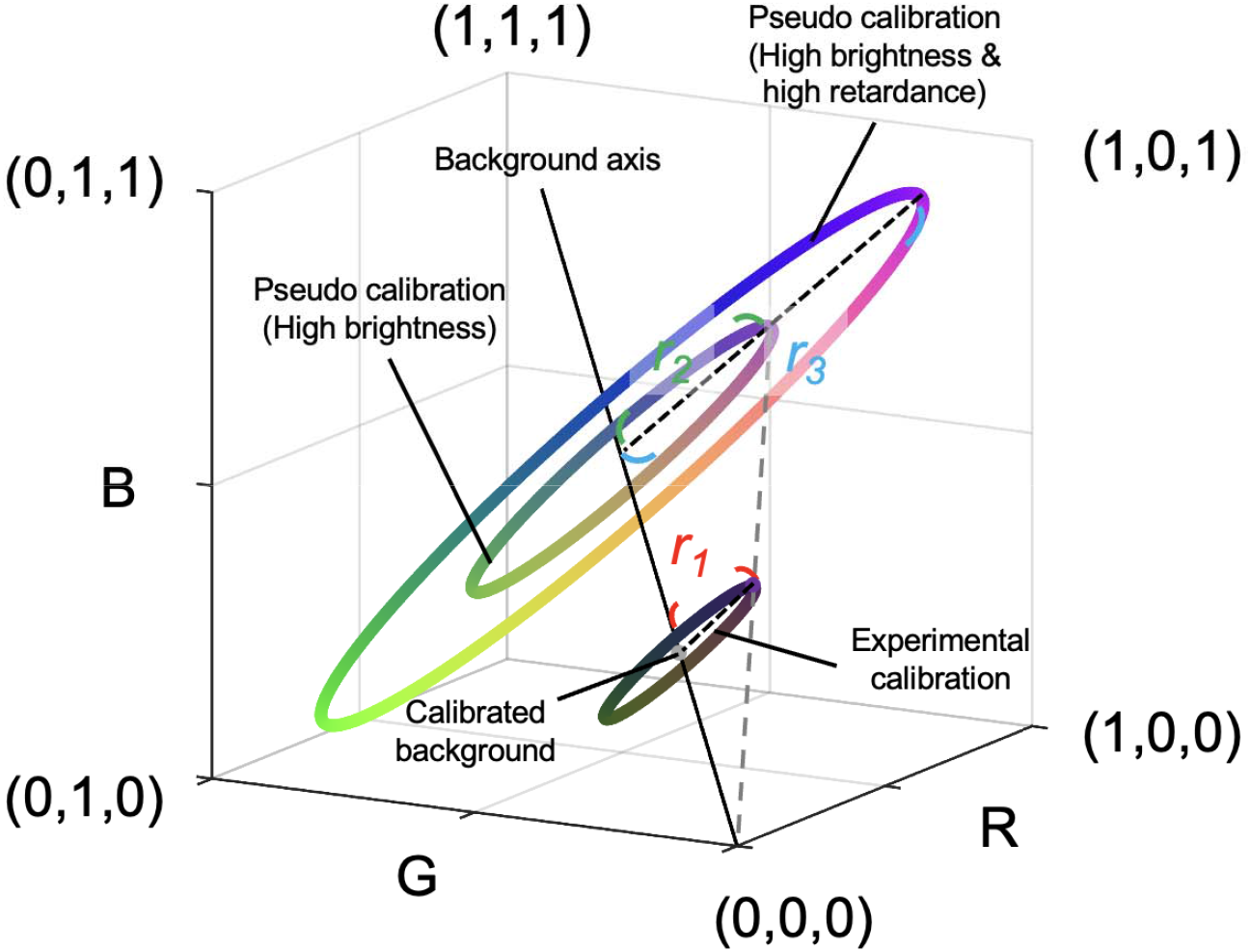
Generation of pseudo calibrations from the experimental calibration. An experimental calibration created a ring with a calibrated background (i.e., central point) and a radius *r*_1_ in the color space. We defined the RGB distance between color on experimental calibration and calibrated background (i.e., radius) as a unit of “energy”, which was related to retardance. Pseudo calibrated rings were generated based on changes in brightness and energy. On one hand, an increase in brightness results in an increase in RGB values in proportion due to exposure time or tissue absorptance. For example, a pseudo calibration ring with a radius *r*_2_ represents a higher exposure time or lower tissue absorption than the experimental calibrated result. Although their radiuses are different, both have the same energy. On the other hand, an increase in energy results in an increase in the radius of the calibrated ring in proportion. For example, a sample generating a calibrated ring with a radius *r*_3_ has higher energy than a sample generating a calibrated ring with a radius *r*_2_. A combination of pseudo and experimental calibrations was used to build interpolating functions for orientation and energy inquiry.

### 2.4 Quantification and visualization

Fiber orientation and energy maps for all images were obtained by searching RGB values for all pixels over the color-angle and color-energy interpolating functions, respectively, obtained from Section 2.3. For visualization, the post-processed image was made from the orientation map of the collagen fibers rendered by a HSV colormap and the brightness was weighted by the energy map.

### 2.5 Imaging collagen architecture and deformation

#### Sample preparation for static imaging

A normal one-year-old sheep eye was obtained from a local abattoir within four hours after death and formalin-fixed for 24 hours at 22 mmHg of intraocular pressure. (Jan et al., 2015a) The muscles, fat, and episcleral tissues were carefully removed. The optic nerve head region was isolated using an 11-mm-diameter trephine and embedded in optimum cutting temperature compound (Tissue-Plus; Fisher Healthcare, TX, USA).

#### Sample preparation for uniaxial stretch testing

An unfixed sheep eye section was prepared as described in detail previously (Jan et al., 2022). Briefly, a normal one-year-old sheep eye was obtained from a local abattoir within four hours after death. The muscles, fat, and episcleral tissues were carefully removed. The optic nerve head region was isolated using an 11-mm-diameter trephine and embedded in optimum cutting temperature compound (Tissue-Plus; Fisher Healthcare, TX, USA). Samples were then snap frozen in liquid nitrogen-cooled isopentane and sectioned coronally at a thickness of 16 μm. OCT was washed with multiple PBS baths. To prevent curling or tears at the clamp points, a tissue section was sandwiched between two pieces of silicone sheet (Medical Grade, 0.005”; BioPlexus, AZ, USA). The sheets also allowed using PBS to maintain tissue hydration without lensing.

#### Static imaging for optic nerve head collagen architecture

Tissue samples were imaged using IPOLπ and IPOL with a 10x strain-free objective (numerical aperture [NA] = 0.3). Due to the limited field of view of the objective, mosaicking was used to image the whole section. The mosaics were obtained with 20% overlap and stitched using Fiji. (Schindelin et al., 2012)

#### Dynamic imaging for sclera while under uniaxial stretching

Each section was mounted to a custom uniaxial stretching device and then stretched on both sides equally and dynamically. The section was imaged using IPOLπ with a 4x strain-free objective (NA = 0.13) to visualize a sclera region. The frame rate was 60 frames per second. Angles within a scleral region were extracted from the initial and three stretching states. Each two states had a 240-frame gap.

## 3. Results

### 3.1 IPOLπ simulation

A simulated RGB color map of IPOLπ presented the interaction with collagen fiber orientation angles from 0 to 180 degrees and retardance from 0 to π radian (**Figure 3a**). Although the colors looked like a continuous rainbow with orientations in the colormap, the relationship of retardance to hue was not a constant and the relationship to brightness was not a strictly increasing function (**Figure 3b**). Therefore, neither hue nor brightness was a suitable parameter for quantifying collagen fiber orientation and retardance, particularly in low-retardance tissues. In the RGB color space, the simulated result appeared as concentric circles, where the radius increased with an increase in retardance (**Figure 3c**). In addition, a non-birefringent sample was located at the central point of the circles. We found that the normalized radius followed the sine function of the retardance, which was equivalent with what we have called normalized “energy” in our previous studies. (Jan et al., 2015a; Jan et al., 2017b) (**Figure 3d**). The explanation of the relationship between the normalized radius and the energy is addressed in more detail in the Discussion. Types of quarter waveplates to build IPOLπ would remarkably impact how the simulated curves within 180-degree orientation in the RGB color space (**Figure 3e**).

**Figure 3.**
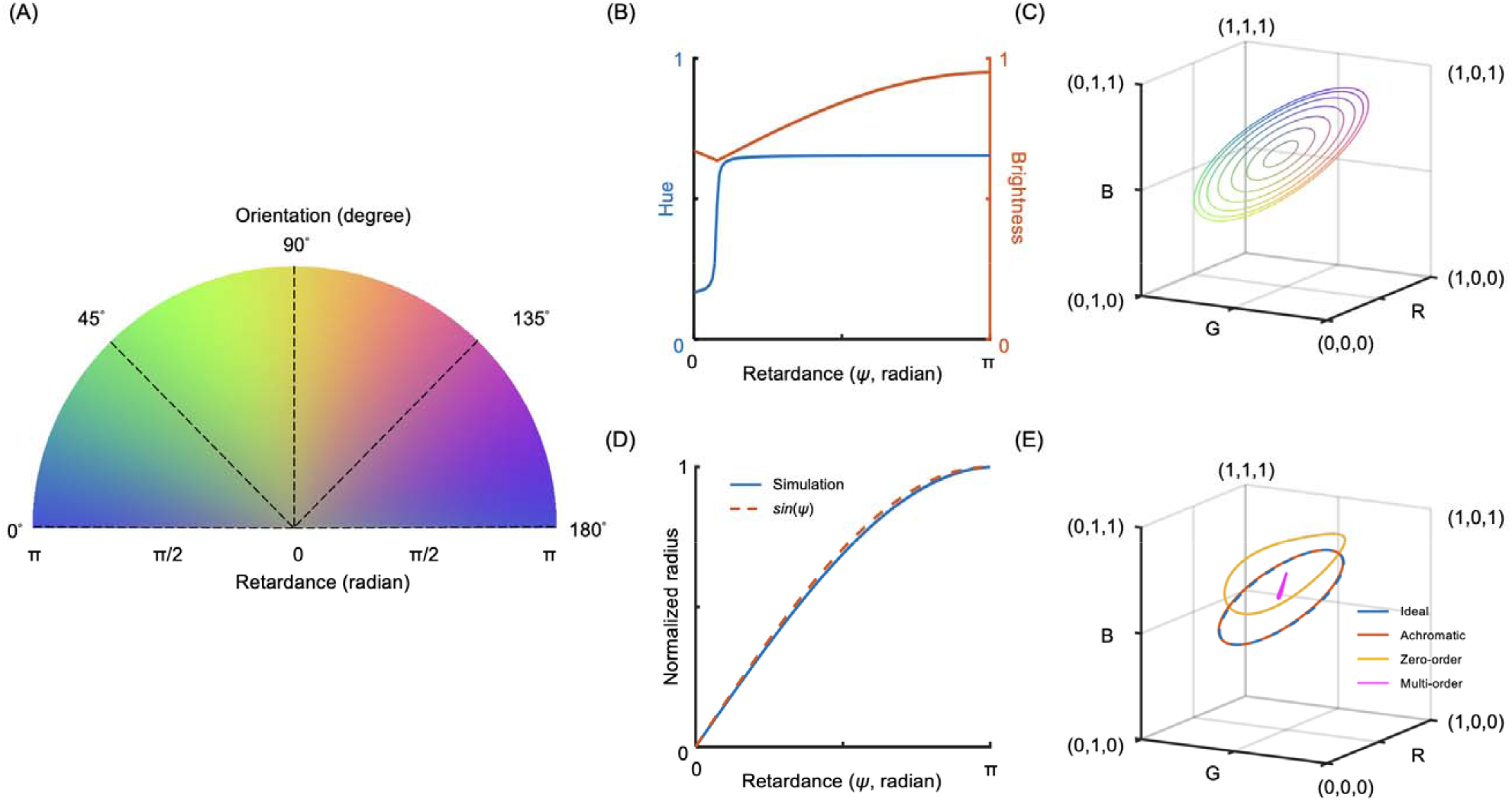
IPOLπ simulation. (a) A simulated RGB color map of IPOLπ based on an achromatic quarter waveplate covering fiber orientation angles from 0 to 180 degrees and retardance from 0 to π radians. (b) Extracting collagen oriented at a 0 degree from (**a**), the relationship between retardance and hue is not a constant and the relationship between retardance and brightness is not a strictly increasing function. (c) Extracting from (a), each simulated ring in the RGB color space corresponds to a collagen fiber with a constant retardance oriented at 0 to 180 degrees. The RGB curves with increased retardance present as concentric circles with increased radiuses, whereas the RGB color for a sample with no retardance is located at the central point of the circle independent of the orientations. (d) The normalized radius obtained from (c) follows the sine function of the retardance, which is equivalent with the normalized “energy”. (e) Each simulated curve corresponds to IPOLπ configured with a different type of quarter waveplate, to acquire a collagen fiber oriented at 0 to 180 degrees. An RGB curve obtained from using an achromatic quarter waveplate is similar to the curve obtained from using an ideal quarter waveplate. A distorted RGB ring obtained from a zero-order waveplate is still colorful, and thus allows building an interpolating function that produces interpolated orientations and energies at inquiry RGBs. Using a multiorder waveplate to build IPOLπ lacks the ability to generate colorful images to visualize collagen microstructure.

### 3.2 RGB color calibration

Images of the chicken tendon section were acquired every 2 degrees from 0 to 180 degrees. The color extracted from the images is presented in the RGB color space with a representative subset of 8 images (**Figure 4a**). The color of the sample acquired by IPOLπ depended on its orientation. All colors formed an enclosing ring in the RGB color space, with the background color located at the center of the ring. A combination of pseudo and experimental calibrations was generated with the corresponding orientations (**Figure 4b**) and energies (**Figure 4c**). These scatter points were used to build two interpolating functions that produced interpolated orientations and energies, respectively, at inquiry RGBs. Note that these interpolating functions are specific to our system. Other systems will potentially vary and therefore require calibration.

**Figure 4.**
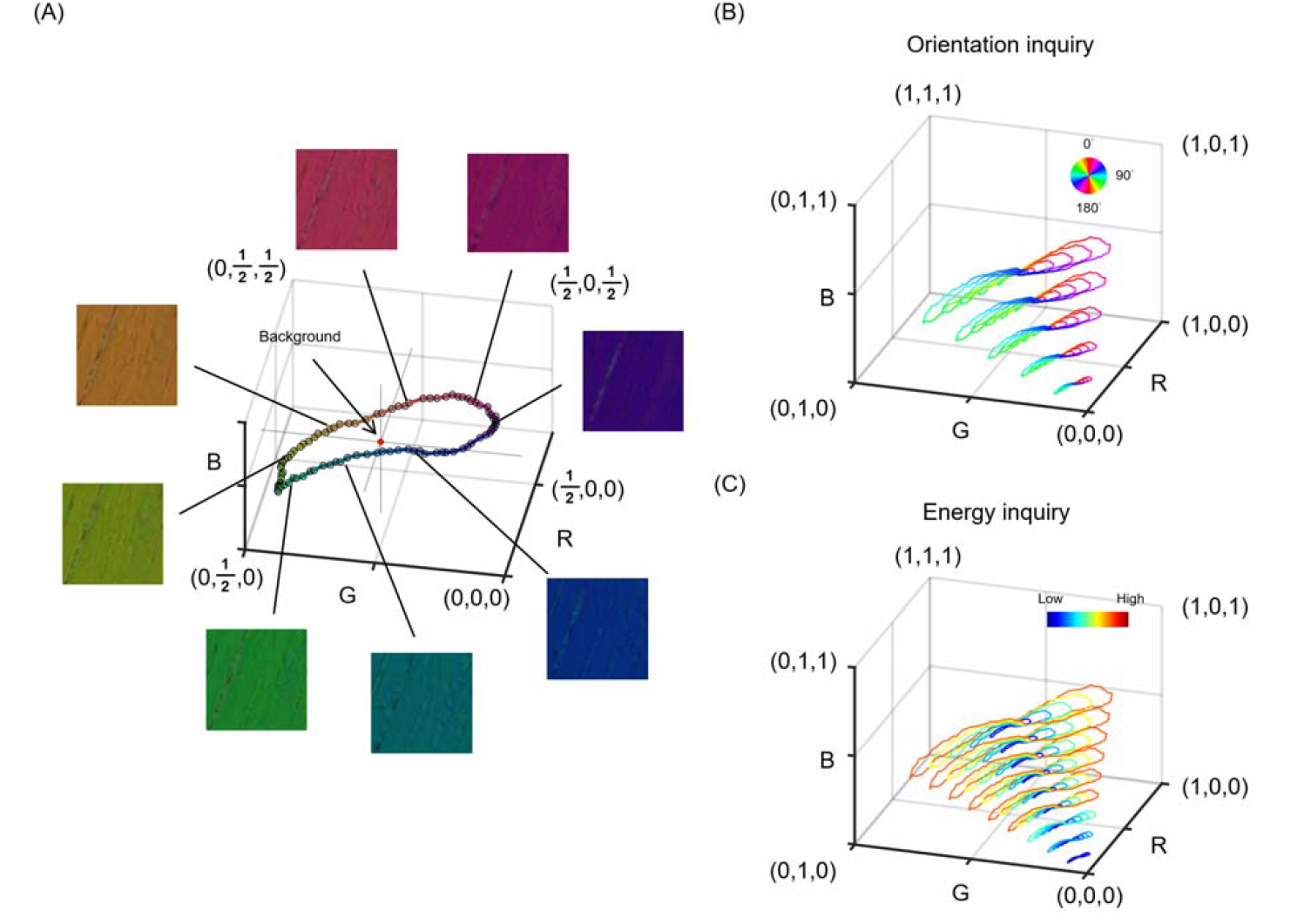
The calibration curve of IPOLπ in the RGB color space. (a) RGB values were obtained from chicken tendon sections at different orientations using IPOLπ. The calibration curve was a ring, where the average of RGB values from all images was the background color (red spot). (b) A combination of pseudo and experimental calibrations was used to build an interpolating function that produced interpolated orientations at inquiry RGBs. Color represents fiber orientation. (c) A combination of pseudo and experimental calibrations was used to build an interpolating function that produced interpolated energies at inquiry RGBs. Color represents energy. Note that the energy in low-brightness pseudo calibrations was weighted by the brightness to avoid noise artifacts.

### 3.3 Collagen architecture and deformation

IPOLπ allowed visualizing the collagen microstructure and orientation in high spatial and angular resolutions (**Figure 5**). The raw image allowed identifying birefringent (e.g., collagen and neural tissues) and non-birefringent (e.g., pigment, and background) components. Collagen and neural tissues are easy to distinguish due to the large differences in birefringence. These are, in part because of their composition, but also because the axons are primarily perpendicular to the section and therefore have lower birefringence than the collagen fibers that are primarily in the plane. (Axer et al., 2001; Yang et al., 2018b) Collagen fibers in the lamina beams and in the scleral canal have clear differences in color, illustrating the strength of IPOLπ having color cycles every 180 degrees compared with conventional IPOL with color cycles every 90 degrees. This help distinguish fibers, even in dense sclera and in complex lamina beams. The post-processed image shows collagen orientation quantitatively. This grey background or region was converted into dark in the energy map. This enhanced the contrast between birefringent and non-birefringent architecture. Figure 6a shows an example IPOLπ image mosaic of a coronal section of sheep optic nerve head. The post-processed image shows orientation and energy information (**Figure 6b**), allowing further quantitative analysis at different scales. At a large scale, IPOLπ allows the calculation, for instance, of regional fiber anisotropy. At a small scale, IPOLπ allows identifying collagen microscale properties such as crimp, or the natural waviness of collagen fibers, discernible in both the PPS and LC regions (**Figure 6c**). IPOLπ can capture dynamic deformations such as uniaxial stretch testing (**Figure 7a**). All post-processing can be conducted after the testing (**Figure 7b**). High spatial and temporal resolutions of IPOLπ enabled the visualization and quantification of the process of load-induced collagen fiber re-orientations (**Figure 7c**).

**Figure 5.**
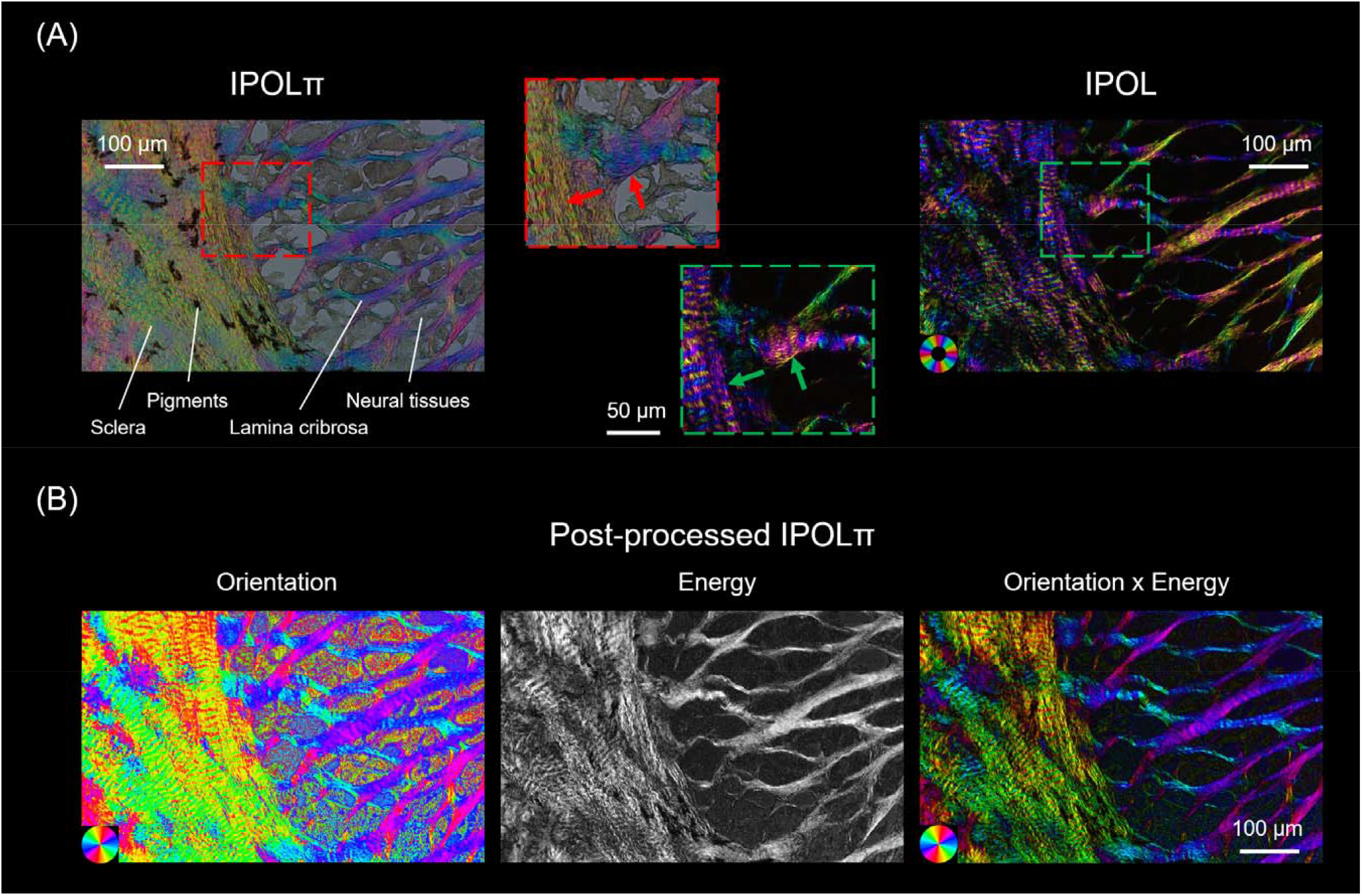
(A) Comparison between IPOLπ (left) and IPOL (right) images as acquired of a section through the lamina cribrosa. The region is at the edge of the scleral canal where the sclera and lamina cribrosa meet. The IPOLπ image has a grey background and the colors low saturation. The dark regions correspond to pigments and the grey textured areas to neural tissues, primarily axons, but also astrocytes and microglia. One of the major differences between IPOLπ and IPOL is the orientation-encoded color cycle. The IPOLπ orientation-encoded color cycle is 180° and thus allows distinguishing the orientations of orthogonal fibers (red arrows) via color. In contrast, the IPOL orientation-encoded color cycle is 90°, which indicates IPOL cannot directly identify the orientations of the orthogonal fibers (green arrows) via color. (B) Post-processed IPOLπ. Orientation and energy maps were extracted from the IPOLπ image using corresponding interpolating functions. For visualization, the “Orientation x Energy” image was obtained from the orientation map of the collagen fibers with brightness scaled by the energy map. The color of the “Orientation x Energy” image had a high contrast and saturation, and regions with low birefringence appear dark. This helps understand the architecture of collagen fibers.

**Figure 6.**
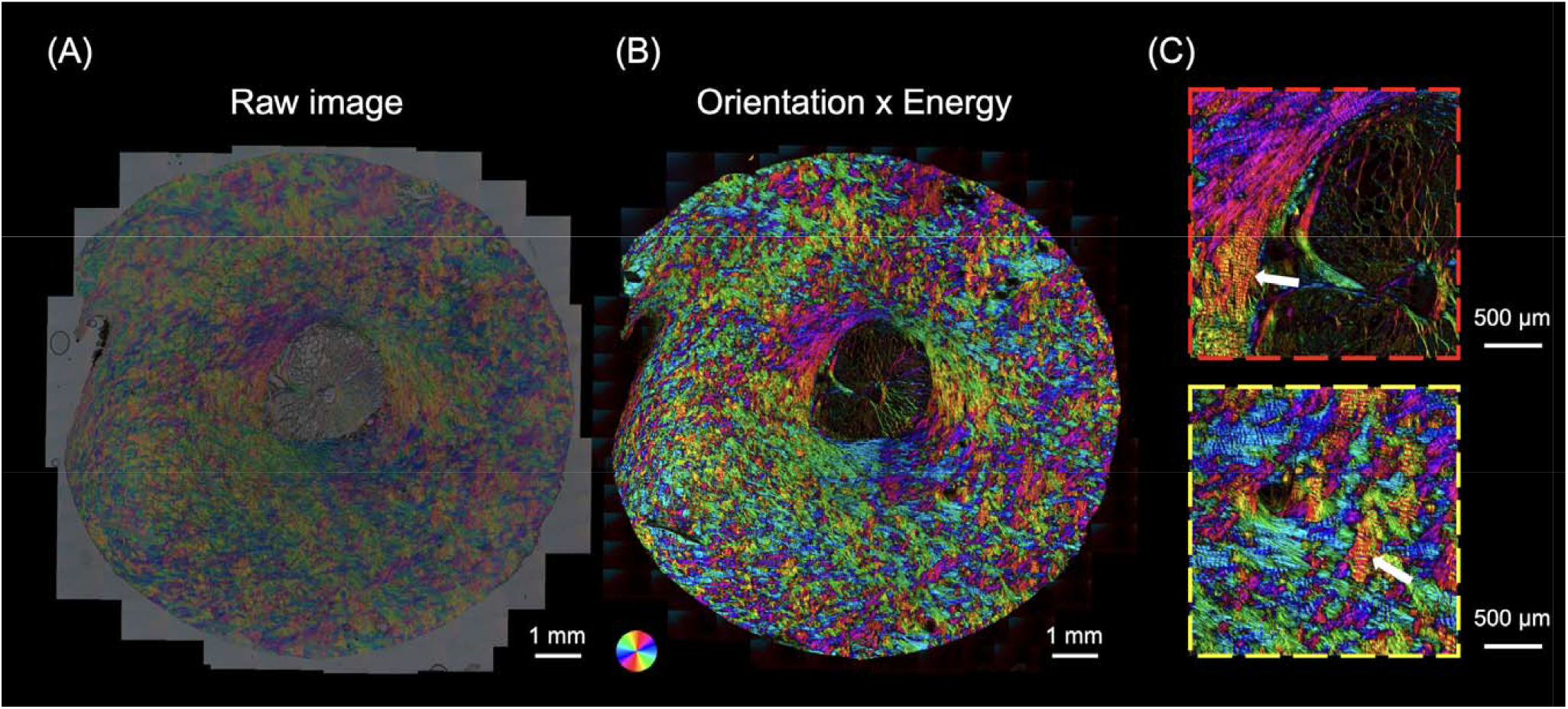
(a) A IPOLπ image acquired as a mosaic of a coronal section from the optic nerve head of a sheep eye. (b) The postprocessed color image was calculated from (a). The color disc on the top left-hand side of the image represents local fiber orientation, and the brightness in the image represents energy. (c) Close-up images from (b) shows interweaving in the peripapillary sclera (red box) and collagen beam networks in the lamina cribrosa (yellow box). The high-resolution image allows identifying crimp (white arrows), an important element of the microstructure for ocular biomechanics.

**Figure 7.**
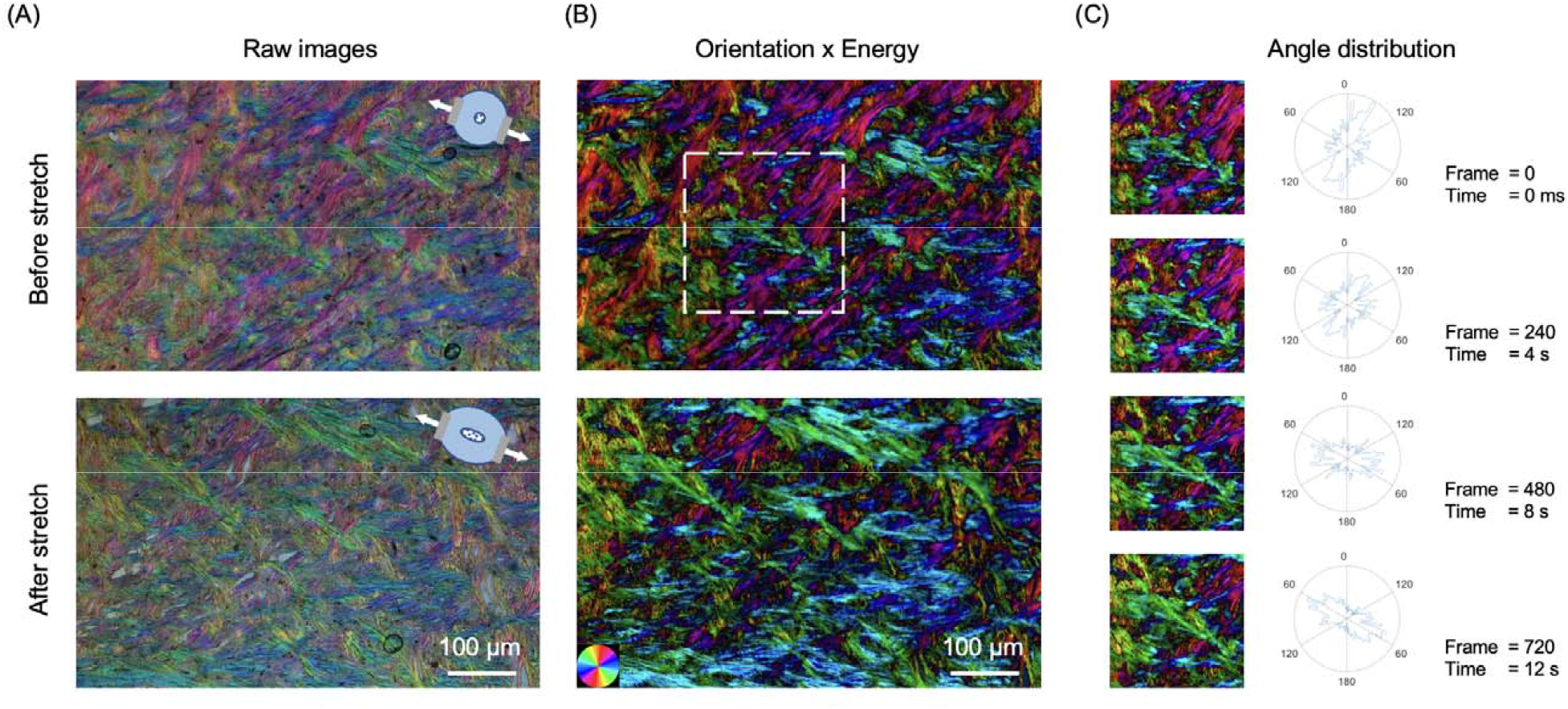
IPOLπ can capture dynamic deformations of ocular tissues. (a) IPOLπ images of an optic nerve head section before stretch and after uniaxial stretch testing. The diagrams on the top right side indicate the stretch directions. (b) The corresponding post-processed color images obtained from the orientation maps masked by the energy maps. (c) Time-sequence images and the corresponding angle distributions from a small interest of region (white box). In the 720^th^ frame, the high frequencies of fiber orientations are close to the stretch directions. The angle distribution plots show the expected change in preferential fiber orientation in the direction of stretch.

## 4. Discussion

Our goal was to demonstrate IPOLπ, a new variation of IPOL in which the orientation-encoding color is cyclic every 180 degrees (π radians). We have shown that IPOLπ produces color images with high spatial, angular and temporal resolutions, suitable for visualization and quantitative analysis of the architecture and dynamics of optic nerve head tissues. Using simulations we built a foundation to convert RGB color into orientation and energy information, and have proved that the orientation-encoded color is cyclic every 180 degrees. The quantitative capability of IPOLπ enables further study on essential biomechanical properties of collagen, such as fiber anisotropy and crimp. We have illustrated the application of IPOLπ high frame rates to show fiber reorientation in uniaxial stretch tests. Altogether, the high spatial and temporal resolutions of IPOLπ enable a deeper insight into ocular structure and biomechanics, and through this on eye physiology and pathology. IPOLπ combines features of conventional PLM and IPOL and that we discuss in detail below. Interestingly, through this work we have also shown that IPOLπ has several convenient properties beyond the color cycle. Further down in the discussion we address these, explaining the strengths and potential applications.

Both IPOLπ and conventional PLM allow distinguishing fiber orientations ranging from 0 to 180 degrees since both system configurations include a circular polarizer that can differentiate the polarization directions of light ranging from 0 to 180 degrees. (Jan et al., 2015a; Kalwani et al., 2013) Images acquired using conventional PLM are monochromatic, and thus the selection of the circular polarizer can be specific to the use of the imaging wavelength. In contrast, in IPOLπ, white light is used to acquire images. Since there is no ideal circular polarization (i.e., perfectly wavelength-independent), the output polarization of each wavelength other than the specific wavelength through the circular polarizer is not ideal circular polarization. The ability of the circular polarizer to minimize the difference in the output polarization of each wavelength affects how the output color changes with collagen orientation, which further affects the capability of distinguishing collagen orientation via color. In Figure 3E, we demonstrate IPOLπ configured with a circular polarizer made by an achromatic or zero-order quarter waveplate can produce a clear color for visualization and quantification of collagen orientation.

Both IPOLπ and conventional PLM allow quantifying the retardance of the collagen structure. Conventional PLM requires images with different polarization states to derive information such as fiber orientation and retardance (Jan et al., 2015a; Mehta et al., 2013). In the four-frame algorithm without extinction setting, the measured retardance δ can be calculated as (Shribak and Oldenbourg, 2003)

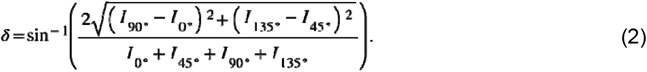

where four intensity measurements, I_0°_, I_45°_, I_90°_, and I_135°_, are taken with linear polarizer analyz**er** set to 0°, 45°, 90°, and 135°, respectively. We previously defined an “energy” for visualization as (Jan et al., 2015a; Jan et al., 2017b)

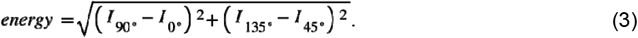

From Equations 2 and 3, we know energy is the sine function of the retardance without normalization. In Figure 3D, we found that the normalized RGB distance (i.e., radius) between the calibrated color and calibrated background color followed the sine function of the retardance. Therefore, it indicates the RGB distance is equivalent with “energy”.

IPOLπ has faster imaging speed and preserves higher resolution than conventional PLM due to the need in PLM of acquiring images with different polarization states. Imaging speed limited by the time required to change the analyzer orientation often precludes conventional PLM from applications on dynamic tissue deformation. (Tower et al., 2002) Several techniques have been demonstrated to improve the imaging speed, such as multiplexing on a single image sensor (Gruev et al., 2010; Kaminsky et al., 2007) and switching polarization states quickly (Keikhosravi et al., 2017). However, these techniques still suffer from post-processing, i.e., image registration and denoising, which takes time and may introduce errors. The interpolation for image registration may introduce errors in orientation and retardance calculations. For large samples, mosaicking is required. Stitching among multiple views also interferes with image registration in a single view, sometimes causing in a large error. Using a camera with built-in multi-analyzer grids on the imaging sensor is a powerful solution for conventional PLM, which acquires images with four polarization states in a snapshot (York et al., 2014). However, having the analyzers integrated, such camera is limited in other applications where it would be preferable to not have the analyzers. In IPOLπ, placing z-cut quartz beside the analyzer allowed diverging the polarization directions of the visible spectrum, generating a colorful light related to collagen fiber microstructure and orientation. IPOLπ only needs a color snapshot. Post-processing is just for color inquiry at a single pixel and thus can be computed in parallel or after imaging. Therefore, IPOLπ preserves the original spatial and temporal resolution offered by the imaging system.

Z-cut quartz induces different polarization rotations within the spectrum, thus affecting the output spectrum and forming the color in IPOLπ. Polarization rotation involves amount and direction. The polarization rotation amount through z-cut quartz depends on the thickness of z-cut quartz and the wavelength of the light. Thicker z-cut quartz increases the difference in polarization rotation within the spectrum. To achieve IPOLπ orientation-encoded color close to “a hue color wheel” (red-yellow-green-blue-red), a suitable thickness of z-cut quartz is critical, which causes enough polarization rotation differences (~180 degrees) in a visible spectrum. Since the thickness of z-cut quartz is mm-level, the effects of parallelism and surface flatness on polarization rotation difference can be ignored. The polarization rotation direction depends on the handedness of z-cut quartz. Therefore, the handedness of z-cut quartz only affects whether the IPOLπ orientation-encoded color is in clockwise or counterclockwise order.

Both IPOLπ and IPOL imaging techniques are based on z-cut quartz to modulate the polarization directions of the visible spectrum, and thus can acquire different polarizations of light via a snapshot. We note three similarities in imaging capability. First, both allow direct visualization of collagen architecture in color. Second, both leverage the full spatial resolution of the microscope-camera system since both use a single snapshot to acquire birefringent information. Third, theoretically, both acquisition speeds are limited only by the camera, thus both are suitable for imaging tissue dynamics and biomechanics.

The configuration of IPOLπ includes a quarter waveplate but not in IPOL, resulting in different imaging features. We highlight three important strengths of imaging capability of IPOLπ compared to the traditional 90-degree IPOL. First, the IPOLπ image can identify the orientations of orthogonal collagen fibers (e.g., laminar beams and collagen fibers in the scleral canal) through colors since the orientation-encoded color in IPOLπ was 180-degree cyclic. In IPOLπ, when the circularly polarized light passes through the birefringent sample, the light becomes elliptically polarized. The analyzer can distinguish the elliptically polarized light with orthogonal axes. In contrast, the orientation-encoded color in IPOL is 90-degree cyclic. IPOL 90-degree cyclic due to its crossed polarizer’s setup. In Malus’s Law, the output intensity of linearly polarized light varies the square of the cosine (i.e., 90-degree cyclic) of the angle between the polarization angle and the transmission axes of the analyzer. Second, acquiring a IPOLπ image only required 1/100 exposure time than acquiring a conventional IPOL image, thus allowing faster imaging speed. The conventional IPOL includes a cross-polarizer design. The design causes a dark background and also causes a low optical transmission rate except for birefringent regions with high retardance. In contrast, a combination of a circular polarizer and an analyzer in IPOLπ does not block the light, and thus the optical transmission rate is high, independent of retardance. Third, IPOLπ allows identifying tissues with high absorption (e.g., pigment and axon) but without birefringence. These non-birefringent tissues appear dark gray. This allows identifying the spatial relationship between collagen and non-birefringent tissues. For example, IPOLπ can visualize the relative displacement and deformation between collagen and axon under stretch testing. For visualization, post-processing for IPOLπ allows for making the non-birefringent tissues invisible. In contrast, no information about these non-birefringent tissues using conventional IPOL since they appear very dark in IPOL.

The IPOLπ system setup is cheaper and simpler than the IPOL’s. IPOL requires two pieces of z-cut quartz, one right-handed and another left-handed. IPOLπ only requires one, which can be of either type. In our experience the left-handed z-cut quartz costs more than triple that of the right-handed quartz. This alone should result in reduced costs. In addition, IPOLπ can be implemented with an imperfectly collimated light source. Polarization rotation through the z-cut quartz is sensitive to the optical path length. In IPOL, the imaging system (e.g., dissecting microscope) requires an extra light source collimator to make sure that a collimated light passes through z-cut quartz in the illumination path. (Lee et al., 2020) This requirement increases the system’s complexity and cost. In IPOLπ, the z-cut quartz can be placed directly into the imaging path. Background artifacts are minimal since the light passing through the objective lens is collimated. We acknowledge that when we set out to develop IPOLπ our motivation was to extend the color cycle to 180 degrees, which was essential to our applications. Identifying the other strengths was somewhat fortuitous. It is not difficult now to imagine situations where these strengths could be the primary motivation for choosing IPOLπ over regular IPOL. An example of the serendipity of science and the importance of research.

We have also shown how to establish quantitative color-angle and color-energy mappings using an experimental calibration. The mapping technique can be applied to any regular color cameras without automatic white balance control and single-color cameras with switching RGB color filters. Note that all optical elements, such as the light source, optical filters and camera parameters may impact the colors. Hence, the calibration is system specific. It is worth noting that adjusting the brightness of the image should be based on the change in exposure time or the use of a neutral density filter. Changing the power of the light source may change the spectrum and thus affect the colors observed in IPOLπ and in turn the mapping functions. Since evaluating the interpolating functions at every color pixel takes time, we produced two 256×256×256 RGB lookup tables for orientation and energy inquiry, respectively, which replaced time-consuming interpolation with an indexing operation. For a 20000×20000 mosaic image, sufficient to include a full section of the posterior pole of the eye at 0.5μm/pixel resolution, the computational time was reduced from 24 minutes to 6 seconds after using the lookup table (1/240^th^ of the time). The approach to quantifying local orientation and energy is powerful and fast, as shown elsewhere in this paper. The color information at each pixel allows us to characterize collagen fiber architecture without tracing individual fibers. Other imaging techniques trace individual fibers through recognizing fiber edges (Cheng et al., 2018; Wu et al., 2003), but these suffer in regions of high fiber density where the edges may be difficult to discern. To avoid this problem it is common to increase the magnification, but this limits the field of view. We acknowledge that the experimental calibration and quantitative method can be improved. Although we demonstrated chicken tendon tissue sections as calibration samples, they are imperfect. Commercial retarders or other synthetic materials may be necessary if finer calibration is necessary. We have illustrated in Figure 3 that the HSV quantitative method is not suitable for IPOLπ, and have demonstrated a quantitative method in the RGB space based on 3D interpolation. Nevertheless, there may exist another color space whose parameters could be simply used for IPOLπ quantification.

It is important to consider the limitations of IPOLπ. First, IPOLπ, as presented in this work, is based on transmitted light illumination, which requires thin samples. For soft tissues this often means cutting them into thin sections, which limits the biomechanics applications. In the future it may be possible to use structured light illumination to extend IPOLπ to work with reflected light, and thus be suitable for imaging thick tissues and potential for in vivo imaging. (Yang et al., 2018a). Second, the diffraction artifacts from fiber edges in IPOLπ lead to a sharply high energy obtained after RGB color inquiry. Therefore, to build a color-energy interpolating function for visualization and avoid artifacts, the energy in low-brightness pseudo calibrations was weighted by the brightness. Third, tissue with selective absorption (e.g., stain) may affect the output color, and thus induce an error in orientation and energy mapping. For example, hematoxylin and eosin (H&E) stain is common in medical diagnosis. (Titford, 2005) Collagen appears strong pink with H&E stain and its orientation color in IPOLπ would be depressed by the stain. Fourth, the colorful light of IPOLπ contains information on all birefringent tissue components, including various types of collagens and non-collagenous components, such as elastin and neural tissue microtubules. (Waxman et al., 2022) Since the birefringence of collagen is substantially larger than that of other birefringent elements in the eye (Inoué and Oldenbourg, 1998; Waxman et al., 2021) we assumed that the majority of the polarized light interaction was from collagen. In addition, we acknowledge that IPOLπ cannot distinguish among various collagen types. The ability to detect birefringence of other components could also be seen as a strength, as it allows applying of IPOLπ to study the structure and biomechanics of neural tissues. This could be useful, for example, to study the optic nerve head of rodents, which have a glial, non-collagenous lamina. (Tamm et al., 2017)

In conclusion, we present IPOLπ as a label-free imaging technique for imaging collagen tissues in the eye with high spatial, angular, and temporal resolutions. IPOLπ optically encodes collagen orientation and retardance in color at each pixel, and thus a color snapshot allows visualization and quantification of collagen architecture. This study provides a novel imaging modality to collagen microstructure and biomechanics in the eye, which could help understand the role of collagen microstructure in eye physiology, aging, and in biomechanics-related diseases.

